# Wild birds as reservoirs of multidrug-resistant enterobacteria in Mulungu, Brazil

**DOI:** 10.1101/2021.11.04.467336

**Authors:** Antonio Jackson Forte Beleza, William Cardoso Maciel, Arianne Silva Carreira, Adson Ribeiro Marques, Carlos Henrique Guedes Nogueira, Neilton Monteiro Pascoal Filho, Bruno Pessoa Lima, Isaac Neto Goes da Silva, Ruben Horn Vasconcelos, Leandro Rodrigues Ribeiro, Régis Siqueira de Castro Teixeira

**Affiliations:** Postgraduate Program in Veterinary Science, Veterinary Faculty, State University of Ceará. Av. Dr. Silas Munguba, 1700, Campus do Itaperi. Zip Code: 60740-903 Fortaleza, Ceará, Brasil; Laboratory of Animal Anatomy and Pathology, Federal University of the Agreste of Pernambuco. Av. Bom Pastor, s/n.° - Boa Vista, Garanhuns – PE, 55292-270; Fazenda Haras Claro, Rod Br 020 Km 381 Bom Princípio, 61600900 Caucaia, CE

## Abstract

Caatinga is a biome unique to Brazil that is degraded by anthropogenic actions, which lead to the loss of biodiversity putting many species at risk of extinction. The Ceará State is located in the Caatinga and has a rich avifauna comprised of 433 species including 13 species that are threatened with extinction, which are found in the Baturité Massif. The aim of this study was to investigate the frequency and diversity of enterobacteria in wild birds and to determine their susceptibility to antimicrobials. Cloacal swab samples were collected from 50 individuals of 28 different species, including the Ceara Gnatheter (*Conopophaga cearae*) and Red-necked Tanager (*Tangara cyanocephala cearensis*), which are classified as vulnerable (VU) by the Brazilian Ministry of the Environment. A total of 55 isolates belonging to 14 different species of Enterobacteriaceae were identified. Among these, *Pantoea agglomerans* and *Escherichia coli* were the most prevalent species with isolation rates of 36% and 26%, respectively. The highest rate of antimicrobial resistance found was to ampicillin (41.8%), followed by nalidixic Acid (36.3%) and amoxicillin associated with clavulanic acid (32.7%). The drugs with the best efficacy were tobramycin (96.4%), ciprofloxacin (92.6%) and tetracycline (90.9%). Multidrug resistance was observed in 23.5% of the analyzed strains. This research provides important information about the composition of the cloacal microbiota of wild birds in Mulungu, Brazil, as well as their health status. In addition, these results demonstrate that they harbor multidrug-resistant strains of Enterobacteriaceae.

## Introduction

It is estimated that the Caatinga biome is home to 548 species of birds, which are distributed in 74 families and represent 28.6% of the total number of species recorded in Brazil [1]. The Ceará State, Brazil, is located in this biome, which has dry landscapes as its main geographic feature. Nonetheless, it presents other striking phytophysiognomies, such as coastal forests, often associated with extensive mangroves, Cerrado fragments, in addition to remnants of Atlantic Forest and Forest Amazon embedded in the semiarid zone [2, 3, 4, 5], as occurs in the enclaves of humid forest occurring in the Baturité Massif [6]. The Baturité Massif has been under strong anthropogenic pressure and since its original occupation it has suffered severe environmental degradation caused by deforestation, fires, introduction of exotic species, dismemberment of sites, predatory hunting and growth of urban centers, which have been important factors for the alteration of the local biota [7]. In addition, it is home to 13 bird species that are classified as threatened in the Red List of the Brazilian Ministry of the Environment [8]. Therefore, the Baturité Massif is a priority area for avian conservation in Northeastern Brazil [9].

The environmental degradation can promote notable negative consequences on wildlife [10]. Environmental pollution is one of the anthropogenic actions that puts the conservation of avifauna at risk, mainly on aspects related to the dissemination of pathogens that are important for animal and public health, such as Salmonella and other enteropathogens [11, 12]. Furthermore, free-ranging birds may come into contact with residues of antibiotics or resistant microorganisms when exposed to contaminants in the environment in which they live [13, 14, 15]. This may affect the health of birds, considering that factors such as ingestion of antibiotics and infection by pathogenic organisms may alter the microbiota of birds [16, 17, 18].

Several studies with free-living birds have shown that they may carry strains of bacteria from the Enterobacteriaceae family with resistance to multiple antimicrobials [19, 20, 21]. Over several decades, antimicrobial resistance has become a global clinical and public health threat against the effective treatment of common infections caused by resistant pathogens, resulting in treatment failure and increased mortality [22]. The development of bacterial resistance can be explained by the natural evolution of microorganisms. However, the widespread and misuse of antibacterial agents in humans and animals has accelerated this process [22]. In recent years, substantial evidence has been provided linking the high presence of antimicrobial-resistant bacteria in the environment with anthropogenic sources [23, 24]. In this context, there is a growing interest in researches involving the environment, including wildlife, in order to better understand the effects of pollution and antimicrobial resistance derived from anthropogenic impacts in ecosystems [25, 26, 27].

Anthropogenic effects on wildlife are poorly investigated, and the extent to which animal populations contribute to the spread of antibiotic resistance is still unknown. Therefore, considering that there are few studies investigating the contact that free-living birds have with multidrug-resistant enterobacteria in the Ceará State, which are generally limited to a few species, greater elucidation is needed [28, 29, 15, 30].

Hence, this study aimed to investigate the presence of enterobacteria in cloacal swab samples of wild birds captured in the city of Mulungu, Ceará, Brazil, and to determine the phenotypic profile of antimicrobial sensitivity of the isolates.

## Material and methods

### Characterization of the Study Area

This study was authorized by the Brazilian Institute for the Environment and Renewable Natural Resources (IBAMA) with SISBIO protocol number 31847-6 and approved by the Ethics Committee for the Use of Animals of the State University of Ceará (Protocol number 4832011/2014).

The study was carried out in the city of Mulungu, Ceará, which is located in the Baturité Massif that contains 16 cities: Aratuba, Baturité, Canindé, Capistrano, Charity, Guaramiranga, Mulungu, Redenção, Pacoti, Palmacia, Acarape, Barreira, Aracoiaba, Guaiúba, Maranguape and Itapiúna. Within the region known as the Baturité Massif, an Environmental Protection Area (APA) is located at an approximate distance of 120 kilometers from the state capital, Fortaleza. This APA presents its highest peak at 1115 meters in altitude and is composed by tropical pluvial subdeciduous forest and pluvio-nebular subevergreen forest (average annual temperatures of 24° to 26° with average annual rainfall of 1,737.5 millimeters and with hot sub-humid and humid tropical conditions) with trees up to 30 meters high, river springs and waterfalls. This region show a marked contrast to the surrounding semiarid backwoods (sertão) in the middle of a hot dry region. It has high anthropogenic activity, such as agriculture, livestock and urban growth, and presents mostly altered vegetation [31, 32, 33].

### Sample collection

The capture of birds was conducted for 3 months (october, november and december 2019) with the aid of 4 mist nets (Ecotone Mist nets - 1030/12-nailon; length: 12 cm; height: 3.2m; mesh: 30×30cm; denier: 110/2; 4 bags, fixed with rods at the ends).

The nets were placed 20 cm above the ground in linear transects in the forest. These were opened at dawn and closed at dusk (7:00 am to 5:00 pm) and were checked every 20 minutes to remove the captured birds.

The species were identified according to the Avis Brasilis field guide [34] and by consulting the list of birds in Brazil made available by the Brazilian Committee for Ornithological Records - CBRO [35].

Biological samples were obtained using sterile cloacal swabs, which were stored in Stuart medium at room temperature, transported and sent within 48 hours to the Ornithological Studies Laboratory, State University of Ceará (LABEO/UECE) for further microbiological processing. After sampling, individuals were marked by clipping a secondary feather from the right wing before being released back to the wild.

### Microbiological procedure

Once at the Ornithological Studies Laboratory (LABEO), samples were transferred from Stuart media to 5 mL of 1% Peptone Water (Kasvi^®^) and were cultured. The incubation conditions were standardized at 37°C/24h for all the steps of the microbiological procedure. Aliquots of 0.5mL were collected from the peptone water samples and transferred to tubes containing Brain-Heart Infusion (Kasvi^®^) (BHI) and Selenite-Cystine (Kasvi^®^) (SC) enrichment broths. In addition, aliquots of 0.05mL were collected and transferred to Rappaport-Vassiliadis broth (Kasvi^®^) (RP). After incubation, a loopful was collected from each broth and streaked on plates containing Brilliant Green agar (Himedia^®^), Salmonella-Shigella agar (Himedia^®^) and MacConkey agar (Kasvi^®^), following incubation. Different colonies were collected from each plate and were inoculated into tubes containing Triple Sugar Iron Agar (Kasvi^®^). To identify the enterobacteria, biochemical tests were used, including SIM Medium (Himedia^®^), lysine-decarboxylase (LIA) (Kasvi^®^), ornithine-decarboxylase (Himedia^®^), methyl red (VM), Voges-Proskauer (VP) (Himedia^®^), Urea (Dynamic Formula^®^), Simmons Citrate Agar (Himedia^®^), Arginine Decarboxylase (Exodus Cientifica^®^), Malonate Broth (Himedia^®^), Lactose (Merck^®^), Sucrose (Dinâmica^®^), Mannitol (Dinâmica^®^), Arabinose (Dinâmica^®^), Raffinose (Dinâmica^®^), Rhamnose (Dinâmica^®^), Sucrose (Dinâmica^®^), Dulcitol (Dynâmica^®^ ^®^), Adonitol (Dinâmica^®^), Inositol (Sigma^®^) and Sorbitol (Sigma^®^) [36].

### Antimicrobial susceptibility profile

The isolates were submitted to anticrobial susceptibility test using the Kirby-Bauer disk diffusion technique [37], and the inhibition zones were compared to the standards established by the Clinical and Laboratory Standards Institute-CLSI [38]. Eleven antimicrobials of 7 pharmacological classes were tested: Quinolones (Nalidixic Acid, 30μg); Fluroquinolones (Ciprofloxacin, 5μg); Aminoglycosides (Gentamicin, 10μg and Tobramycin, 10μg); Tetracyclines (Tetracycline, 30μg); Macrolides (Azithromycin, 15μg); Sulfonamides (Sulfamethoxazole + Trimethoprim, 25 μg); Beta-lactams (Penicillin: Ampicillin, 10 μg and Amoxicillin + Clavulanic Acid 10 μg, Cephalosporins: Ceftriaxone, 30 μg and Carbapenems: Meropenem 10μg); (All antimicrobials from Oxoid Ltd., Cambridge, UK confirm brand*). Isolates that expressed resistance or intermediate phenotypes were interpreted as resistant. Bacteria were considered resistant to multiple drugs (RMD) when resistance occurred to at least three classes of antibiotics [39]. The *Escherichia coli* ATCC 25922 strain was used as a control sample. To perform the test, isolates were cultured in tubes containing 5 mL of Brain-heart Infusion broth (BHI), and placed in bacteriological incubator for 24 hours at 37°C. Subsequently, aliquots of the broth were seeded onto MacConkey agar plates and again incubated. Afterwards, two to three colonies were selected and placed in 5mL tubes of saline solution. Then, a swab was moistened in the turbid saline solution (which contained a turbidity of 0.5 according to the Mcfarland Nephelometric scale) and streaked on the surface of a plate containing Mueller-Hinton agar (Kasvi^®^), to which antimicrobial discs were placed. After the plates were incubated at 37°C for 24h, the inhibition zones were measured and results were interpreted as sensitive or resistant.

## Results

During the study, a total of 50 birds of 28 different species distributed in 13 families were captured (Table 1). The most frequent species was Pectoral Sparrow (*Arremon taciturnus*), in a total of 5 individuals, followed by the occurrence of Yellow-bellied Elaenia (*Elaenia flavogaster*) and Ruddy Ground-Dove (*Columbina talpacoti*), both with 4 individuals. Two rare species classified as vulnerable (VU) were also collected, which were the Ceara Gnateater (*Conopophaga cearae*) and the Red-necked Tanager (*Tangara cyanocephala cearensis*) [8].

**Table 1.** List of bird species captured in the city of Mulungu, Ceará, Brazil.

| Family | Common and scientific name | Captured animals |
| --- | --- | --- |
| Thraupidae | Red-necked Tanager (*Tangara cyanocephala cearensis*) | 2 |
|  | Burnished-buff Tanager (*Stilpnia cayana*) | 1 |
|  | Bananaquit (*Coereba flaveola*) | 2 |
|  | Palm Tanager (*Thraupis palmarum*) | 3 |
|  | Guira Tanager (*Hemithraupis guira*) | 1 |
|  | Orange-headed Tanager (*Thlypopsis sordida*) | 1 |
| Tyrannidae | Gray Elaenia (*Myiopagis caniceps*) | 2 |
|  | Mouse-colored Tyrannulet (*Phaeomyias murina*) | 1 |
|  | Short-crested Flycatcher (*Myiarchus ferox*) | 1 |
|  | Yellow-bellied Elaenia (*Elaenia flavogaster*) | 4 |
| Columbidae | Gray-fronted Dove (*Leptotila rufaxilla*) | 2 |
|  | Ruddy Ground-Dove (*Columbina talpacoti*) | 4 |
| Trochilidae | Ruby-topaz Hummingbird (*Chrysolampis mosquitus*) | 1 |
|  | Fork-tailed Woodnymph (*Thalurania furcata*) | 2 |
|  | Rufous-breasted Hermit (*Glaucis hirsutus*) | 3 |
| Passerellidae | Pectoral Sparrow (*Arremon taciturnus*) | 5 |
| Dendrocolaptidae | Straight-billed Woodcreeper (*Dendroplex picus*) | 2 |
|  | Planalto Woodcreeper (*Dendrocolaptes platyrostris*) | 1 |
|  | Lafresnaye's Woodcreeper (*Xiphorhynchus guttatoides eytoni*) | 1 |
|  | Red-billed Scythebill (*Campylorhamphus trochilirostris*) | 1 |
| Turdidae | Pale-breasted Thrush (*Turdus leucomelas*) | 1 |
|  | Rufous-bellied Thrush (*Turdus rufiventris*) | 2 |
| Furnariidae | Pale-legged Hornero (*Furnarius leucopus*) | 2 |
| Picidae | Green-barred Woodpecker (*Colaptes melanochloros*) | 1 |
| Conopophagidae | Ceara Gnateater (*Conopophaga cearae*) | 1 |
| Icteridae | Variable Oriole (*Icterus pyrrhopterus*) | 1 |
| Thamnophilidae | Great Antshrike (*Taraba major*) | 1 |
| Hirundinidae | Southern Rough-winged Swallow (*Stelgidopteryx ruficollis*) | 1 |
| Total | 28 | 50 |

A total of 55 strains distributed in 14 different bacterial species were detected in the analyzed samples. The prevalence of positive birds for at least one bacterial species was 52.0%. *Pantoea agglomerans* and *Escherichia coli* were the most prevalent ones, occurring in 36.0% (18/50) and 26% (13/50) of the investigated birds. *Serratia rubidaea* was the third most isolated bacterial species, followed by *Hafnia alvei*, which presentedisolation rates of 14.0% (7/50) and 10.0% (5/50), respectively (Table 2).

**Table 2-.** Absolute and relative frequencies of enterobacteria per bird family in cloacal samples of wild birds captured from October to December 2019 in the city of Mulungu, Ceará, Brazil.

| Isolated<br>Enterobacteriaceae | Total<br>number<br>of birds<br>(n=50) | Absolute and relative frequencies |  |  |  |  |  |  |  |  |  |  |  |  |
| --- | --- | --- | --- | --- | --- | --- | --- | --- | --- | --- | --- | --- | --- | --- |
|  |  | THR<br>(n=10) | TYR<br>(n=8) | COL<br>(n=6) | TRO<br>(n=6) | PAS<br>(n=5) | DEN<br>(n=5) | TUR<br>(n=3) | FUR<br>(n=2) | PIC<br>(n=1) | CON<br>(n=1) | ICT<br>(n=1) | THA<br>(n=1) | HIR<br>(n=1) |
| *Pantoea agglomerans* | 18 (36.0%) | 2 (20.0%) | 1 (25.5%) | 2 (33.3%) | 1 (16.6%) | 3 (60.0%) | 3 (60.0%) | 1 (33.3%) | 1 (50.0%) | 1 (100.0%) | 1 (100.0%) | 1 (100.0%) | 1 (100.0%) | 0 (0%) |
| *Escherichia coli* | 13 (26.0%) | 1 (10.0%) | 1 (25.5%) | 3 (50.0%) | 0 (0%) | 2 (40.0%) | 3 (60.0%) | 2 (66.6%) | 1 (50.0%) | 0 (0%) | 0 (0%) | 0 (0%) | 0 (0%) | 0 (0%) |
| *Serratia rubidaea* | 7 (14.0%) | 0 (0%) | 1 (25.5%) | 2 (33.3%) | 0 (0%) | 1 (20.0%) | 1 (20.0%) | 1 (33.3%) | 0 (0%) | 1 (100.0%) | 0 (0%) | 0 (0%) | 0 (0%) | 0 (0%) |
| *Hafnia alvei* | 5 (10.0%) | 1 (10.0%) | 0 (0%) | 0 (0%) | 0 (0%) | 1 (20.0%) | 2 (40.0%) | 1 (33.3%) | 0 (0%) | 0 (0%) | 0 (0%) | 0 (0%) | 0 (0%) | 0 (0%) |
| *Enterobacter gergoviae* | 2 (4.0%) | 0 (0%) | 0 (0%) | 0 (0%) | 0 (0%) | 1 (20.0%) | 0 (0%) | 1 (33.3%) | 0 (0%) | 0 (0%) | 0 (0%) | 0 (0%) | 0 (0%) | 0 (0%) |
| *Edwardsiella tarda* | 2 (4.0%) | 0 (0%) | 0 (0%) | 0 (0%) | 0 (0%) | 0 (0%) | 1 (20.0%) | 0 (0%) | 0 (0%) | 0 (0%) | 0 (0%) | 0 (0%) | 1 (100.0%) | 0 (0%) |
| *Klebsiella pneumoniae* | 1 (2.0%) | 0 (0%) | 0 (0%) | 0 (0%) | 0 (0%) | 1 (20.0%) | 0 (0%) | 0 (0%) | 0 (0%) | 0 (0%) | 0 (0%) | 0 (0%) | 0 (0%) | 0 (0%) |
| *Proteus vulgaris* | 1 (2.0%) | 0 (0%) | 0 (0%) | 0 (0%) | 0 (0%) | 0 (0%) | 0 (0%) | 1 (33.3%) | 0 (0%) | 0 (0%) | 0 (0%) | 0 (0%) | 0 (0%) | 0 (0%) |
| *Proteus mirabilis* | 1 (2.0%) | 0 (0%) | 0 (0%) | 0 (0%) | 0 (0%) | 0 (0%) | 0 (0%) | 0 (0%) | 0 (0%) | 0 (0%) | 1 (100.0%) | 0 (0%) | 0 (0%) | 0 (0%) |
| *Cronobacter sakazakii* | 1 (2.0%) | 0 (0%) | 0 (0%) | 1 (16.6%) | 0 (0%) | 0 (0%) | 0 (0%) | 0 (0%) | 0 (0%) | 0 (0%) | 0 (0%) | 0 (0%) | 0 (0%) | 0 (0%) |
| *Enterobacter cloacae* | 1 (2.0%) | 0 (0%) | 0 (0%) | 0 (0%) | 0 (0%) | 0 (0%) | 1 (20.0%) | 0 (0%) | 0 (0%) | 0 (0%) | 0 (0%) | 0 (0%) | 0 (0%) | 0 (0%) |
| *Arizona* spp | 1 (2.0%) | 0 (0%) | 0 (0%) | 0 (0%) | 0 (0%) | 0 (0%) | 1 (20.0%) | 0 (0%) | 0 (0%) | 0 (0%) | 0 (0%) | 0 (0%) | 0 (0%) | 0 (0%) |
| *Yersinia enterocolitica* | 1 (2.0%) | 0 (0%) | 0 (0%) | 0 (0%) | 0 (0%) | 0 (0%) | 1 (20.0%) | 0 (0%) | 0 (0%) | 0 (0%) | 0 (0%) | 0 (0%) | 0 (0%) | 0 (0%) |
| *Shigella* spp | 1 (2.0%) | 0 (0%) | 0 (0%) | 0 (0%) | 0 (0%) | 0 (0%) | 0 (0%) | 0 (0%) | 1 (50.0%) | 0 (0%) | 0 (0%) | 0 (0%) | 0 (0%) | 0 (0%) |
| Positive samples | 26 (52.0%) | 2 (20.0%) | 2 (25.0%) | 4 (66.6%) | 1 (16.6%) | 4 (80.0%) | 5 (100.0%) | 3 (100.0%) | 1 (50.0%) | 1 (100.0%) | 1 (100.0%) | 1 (100.0%) | 1 (100.0%) | 0 (0%) |

THR= Thraupidae, TYR= Tyrannidae, COL=Columbidae, TRO=Trochilidae, PAS= Passerellidae, DEN=Dendrocolaptidae, TUR=Turdidae, FUR=Furnariidae, PIC=Picidae, CON= Conopophagidae, ICT= Icteridae, THA= Thamnophilidae, HIR= Hirundinidae

Negative samples were obtained from birds of the Hirundinidae family. On the other hand, all species of the Dendrocolaptidae, Turdidae, Picidae, Conopophagidae, Icteridae and Thamnophilidae family had at least one bacterial isolate. In the Trochilidae family, there was only one (1/6 species) bird that presented bacterial growth, which was a Rufous-breasted Hermit (*Glaucis hirsutus*) that was positive for *Pantoea agglomerans.* Another family with a low number of positive birds was Thraupidae that presented only two birds positive for enterobacteria (2/10 species). An Orange-headed Tanager (*Thlypopsis sordida*) was positive for *Hafnia alvei* and *Pantoea agglomerans*, whereas a Bananaquit (*Coereba flaveola*) was positive for *Pantoea agglomerans* and *Escherichia coli*. The Tyrannidae family had the same number of positive birds (2/8 species), two individuals of Yellow-bellied Elaenia (*Elaenia flavogaster*) from which *Pantoea agglomerans* and *Serratia rubidaea* were isolated from one sample and *Escherichia coli* was isolated from the other. Lafresnaye’s Woodcreeper (*Xiphorhynchus guttatoides eytoni*) was the species with the highest number of isolated enterobacteria (*Enterobacter cloacae*, *Serratia rubidaea, Escherichia coli, Edwardsiella tarda, Hafnia alvei* and *Arizona* spp). The Red-necked Tanager species (*Tangara cyanocephala* subsp. *cearensis*) classified as vulnerable (VU) had no isolates, while the species Ceara Gnateater (*Conopophaga cearae*) classified as vulnerable (VU) was positive for *Proteus mirabilis* and *Pantoea agglomerans* (Table 3).

**Table 3.**
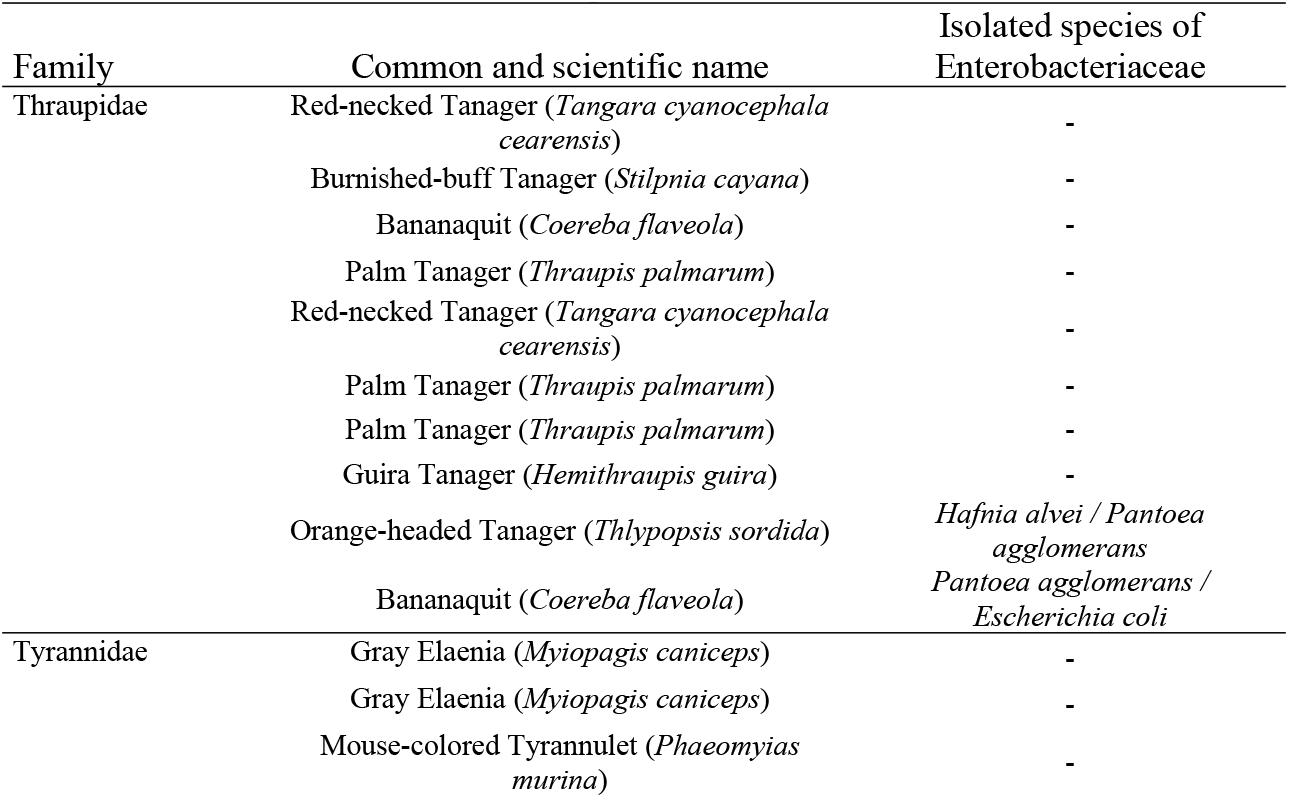
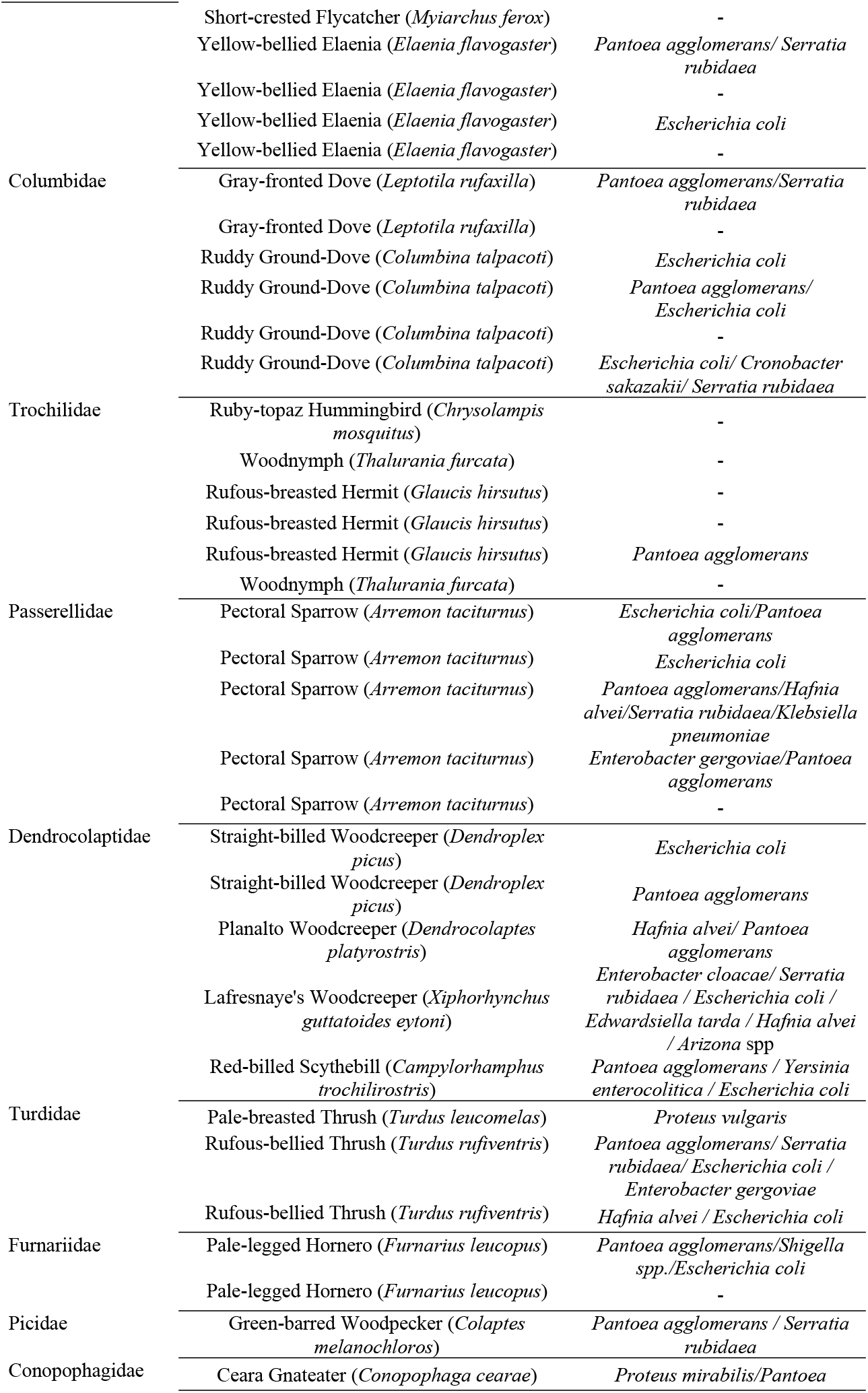
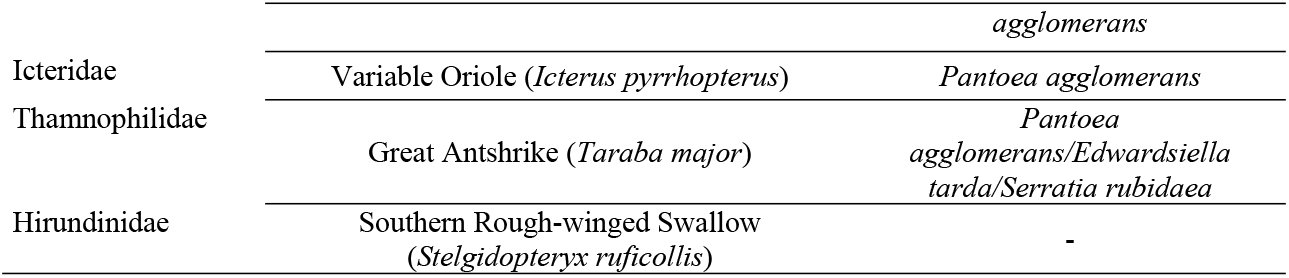
Bacterial species isolated from free-living wild birds captured in the city of Mulungu, Ceará, Brazil.

Considering the total of evaluated strains, the highest rate of antimicrobial resistance occurred to ampicillin 47.3% (26/55). Even after the exclusion of intrinsic resistance (*Klebsiella pneumonia* and *Hafnia alvei*), the rate of 41.8 % (23 strains) was still the highest result. After excluding cases of intrinsic resistance (*Hafnia alvei*), nalidixic acid with a rate of 36.3% (20/55) and amoxicillin associated with Clavulanic Acid with 32.7% (18/55), were the second and third antimicrobials with the highest resistance rates. Tobramycin, ciprofloxacin and tetracycline were the drugs that showed the best efficacy with rates of 96.4%, 92.6% and 90.9% respectively. Meropenem and Gentamicin also performed well (85.5% and 81.8% efficacy, respectively). Considering *E. coli*, the highest resistance rate was also detected to ampicillin, 53.8% (7/13). In contrast, all strains were sensitive to Ciprofloxacin. Regarding the *Cronobacter sakazakii* and *Enterobacter cloacae* strains, both showed resistance only to Nalidixic Acid (Table 4).

**Table 4.** Absolute and relative frequencies of antimicrobial-resistant enterobacteria isolated from cloacal swab samples of wild birds captured from October to December 2019 in the city of Mulungu, Ceará, Brazil.

| Enterobacteriaceae | GEN | AZI | TOB | AMP | CEF | AMO+AC. CLA | CIP | TET | SUL | AC. NAL | MER |
| --- | --- | --- | --- | --- | --- | --- | --- | --- | --- | --- | --- |
|  | n (%) | n (%) | n (%) | n (%) | n (%) | n (%) | n (%) | n (%) | n (%) | n (%) | n (%) |
| *Pantoea agglomerans*, n=18 | 4 (22.2%) | 5 (27.7%)* | - | 7 (38.9%) | 3 (16.6%) | 9 (50%) | 1 (5.5%) | - | - | 7 (38.9%) | 2 (11.1%) |
| *Escherichia coli*, n=13 | 1 (7.7%) | 2 (15.4%)* | 1 (7.7%) | 7 (53.8%) | 2 (15.4%) | 2 (15.4%) | - | 1 (7.7%) | 2 (15.4%) | 2 (15.4%) | 1 (7.7%) |
| *Serratia rubidae*, n=7 | 2 (28.6%) | 2 (28.6%)* | - | 4 (57.1%) | 2 (28.6%) | 1 (14.3%) | 1 (14.3%) | 1 (14.3%) | - | 3 (42.8%) | 2 (28.6%) |
| *Hafnia alvei*, n=5 | - | 2 (40.0%)* | - | 2 (40%)* | 1 (20%) | 2 (40%)* | - | 1 (20%) | 2 (40%) | 2 (40%) | - |
| *Enterobacter gergoviae*, n=2 | 1 (50%) | - | - | 1 (50%) | - | 1 (50%) | - | - | - | - | - |
| *Edwardsiella tarda*, n=2 | - | 2 (100%)* | - | 1 (50%) | - | 2 (100%) | 1 (50%) | - | - | - | - |
| *Klebsiella pneumonia*, n=1 | 1 (100%) | - | - | 1 (100%)* | 1 (100%) | 1 (100%) | 1 (100%) | - | - | 1 (100%) | 1 (100%) |
| *Proteus vulgaris*, n=1 | - | 1 (100%)* | - | - | 1 (100%) | 1 (100%) | - | 1 (100%)* | - | 1 (100%) | 1 (100%) |
| *Proteus mirabilis*, n=1 | - | 1 (100%)* | - | - | 1 (100%) | - | - | - | - | 1 (100%) | - |
| *Cronobacter sakazakii*, n=1 | - | - | - | - | - | - | - | - | - | 1 (100%) | - |
| *Enterobacter cloacae*, n=1 | - | - | - | - | - | - | - | - | - | 1 (100%) | - |
| *Arizona* spp, n=1 | - | - | - | 1 (100%) | - | 1 (100%) | - | - | - | - | - |
| *Yersinia enterocolitica*, n=1 | - | 1 (100%)* | - | 1 (100%) | - | - | - | - | - | - | - |
| *Shigella* spp, n=1 | 1 (100%) | 1 (100%) | 1 (100%) | 1 (100%) | - | - | 1 (100%) | 1 (100%) | 1 (100%) | 1 (100%) | 1 (100%) |
| Acquired resistance | 10 (18.2%) | 1 (1.8) | 2 (3.6%) | 23 (41.8%) | 11 (20%) | 18 (32.7%) | 5 (9.1%) | 4 (7.8) | 5 (9.1%) | 20 (36.3%) | 8 (14.5) |
| Total resistance (acquired + intrinsic) | 10 (18.2%) | 17 (30.9)* | 2 (3.6%) | 26 (47.3%)* | 11 (20%) | 20 (36.3%) | 5 (9.1%) | 5 (9.1%)* | 5 (9.1%) | 20 (36.3%) | 8 (14.5) |

GEN-Gentamicin; AZI-Azithromycin; TOB-Tobramycin; AMP-Ampicillin; CEF-Ceftriaxone; AMO+AC.CLA- Amoxicillin associated with Clavulanic Acid; CIP-Ciprofloxacin; TET-Tetracycline; SUL- Sulfonamide; AC. NAL- Nalidixic Acid; MER- Meropenem; \*- Intrinsic Resistance

Among the birds classified as vulnerable (VU) by the list of the Official National List of Fauna Species Endangered with Extinction [8], it was not possible to detect resistance in samples collected from the Red-necked Tanager (*Tangara cyanocephala cearensis*), since there was no isolation of any bacteria. However, the *Proteus mirabilis* strain that was isolated from a Ceara Gnateater (*Conopophaga cearae*) was resistant to azithromycin, ceftriaxone and nalidixic acid. In addition, the *Pantoea agglomerans* strain that was isolated from the same individual was resistant to eight out of twelve tested antibiotics (gentamicin, nalidixic acid, ceftriaxone, amoxicillin + clavulanic acid, ciprofloxacin, ampicillin, ciprofloxacin and meropenem), which correspond to five of the seven antimicrobial classes.

As expected, when considering the bacterial species that have intrinsic resistance mechanisms, resistance to at least one of the tested antimicrobials was observed in all of the strains. However, when considering only acquired resistance, 10 isolates (18.2%) were sensitive to all of the investigated drugs. Multidrug resistance (acquired cases) was observed in 13 isolates (23.5%), and three strains were resistant to seven antibiotics. From the total of 13 *Escherichia coli* strains, 2/13 (7.7%) presented multidrug resistance and 4/13 (30.8%) of the strains were sensitive to all of the studied antimicrobials (Table 5).

**Table 5.** Absolute and relative frequencies of resistance to multiple drugs of Enterobacteriaceae strains isolated from cloacal swabs from free-living birds captured in the city of Mulungu, Ceará, Brazil.

| Number of antibiotic classes | Frequency of resistant *Escherichia coli* (%) | Resistant enterobacteria |  |
| --- | --- | --- | --- |
|  |  | Only acquired resistance (%) | Total resistance (intrinsic + acquired) (%) |
| 0 | 4 (30.8%) | 10 (18.2%) | - |
| 1 | 5 (38.4%) | 20 (36.4%) | 10 (18.2%) |
| 2 | 2 (15.4%) | 12 (21.9%) | 22 (40%) |
| 3 | 1 (7.7%) | 5 (9.1%) | 13 (23.7%) |
| 4 | - | 2 (3.6%) | 3 (5.4%) |
| 5 | 1 (7.7%) | 3 (5.4%) | 3 (5.4%) |
| 6 | - | - | 1 (1.9%) |
| 7 | - | 3 (5.4%) | 3 (5.4%) |

## Discussion

In this study, more than half of the samples were positive to some of the investigated enterobacteria. Despite the isolation of fourteen different species of bacteria, birds were not necessarily suffering from any pathological condition. In addition to the low frequency of isolation, some of these microorganisms may occur naturally in these birds, considering that these strains have been isolated previously from healthy birds either in the wild or in cages [40, 41, 42, 43, 44, 45, 30]. Species of the Enterobacteriaceae family, *Escherichia coli* in particular, do not belong to the intestinal microbiota of granivorous pet birds, because feed composed exclusively of seeds has been shown to provoke an inhibitory effect of this bacterial species [46]. In this sense, the detection of Enterobacteriaceae in cloacal samples of granivorous birds should be observed with caution, as it suggests favorable conditions for the development of potential pathologies [47]. However, it is important to highlight that the bird species that were captured in this study have an omnivorous diet. This may explain the natural presence of enterobacteria, since the occurrence of these microorganisms in the digestive tract is influenced by the composition of their nutrition [46]. Other factors may also have influenced the isolation of these bacteria, such as direct or indirect contact with domestic animals, as well as environmental contamination by human action. In the natural environment in which they live, several species of enterobacteria also occur, as they are ubiquitous, in soil or water. Thus, Enterobacteriaceae have been isolated mainly from omnivorous, piscivorous and healthy carnivore birds [46, 48, 49, 50, 51].

The most prevalent bacterial species isolated from birds in this study was *Pantoea agglomerans*. This microorganism can rarely cause infections, while it normally acts as a commensal species that colonizes the normal intestinal microbiota. Biodiversity studies report the isolation of *Pantoea agglomerans* in the microbiota of several plants and insects [52, 53], which serve as a food source for several of the bird species that were captured. *Escherichia coli* was also among the most isolated microorganisms (23.2%) and is likewise an ubiquitous organism, which can be found in soil, water and vegetation [54, 55]. Although its presence does not necessarily mean a sign of illness, on some occasions, such as when they acquire virulence genes, this bacterium can cause pathological problems in humans and animals, including birds [41].

The prevalence of *E. coli* in studies involving free-ranging birds is quite varied. Saviolli [56] describes the presence of the microorganism in 60.0% of samples from Magnificent Frigatebird (*Fregata magnificens*) from the coast of the State of São Paulo. Vilela et al. [57] investigated this microorganism in fecal samples of House Sparrows (*Passer domesticus*) that lived around farms in the State of Pernambuco and found lower percentages (13.2%). Callaway et al. [58] analyzed cloacal swab samples from 376 migratory birds, which included Brown-Headed Cowbird (*Molothrus ater*), Common Grackle (*Quiscalus quiscula*) and Cattle Egret (*Bubulcus ibis*), and found even lower rates, 3.7% (14/376).

*Serratia rubidea* was the third most isolated species of bacteria. It is considered an important human pathogen as a common agent of nosocomial infections mainly of the urinary tract [59]. Diseases caused by *Serratia* in birds are uncommon but can occur mostly in an opportunistic manner in immunocompromised birds due to stress in captivity, inappropriate weather conditions, parasitic diseases, among other causes [60, 61]. Free-living birds can either acquire this microorganism from the contaminated environment in which they live and may act as disseminators. Spena et al. [62] isolated *Serratia rubidea* from oral swab samples of Eurasian Thick-knee (*Burhinus oedicnemus*) and associated this finding with a diet composed of invertebrates found in the feces of ruminants. This bacterial species has also been reported to be isolated from lake waters in Poland, which was occupied by Great Cormorant (*Phalacrocorax carbo*). Researchers have associated this finding with the leaching of feces and excreta during rains leading this and other species of enterobacteria into the lake [63].

The other enterobacteria that occurred less frequently in the analyzed samples can also occasionally cause damage to health and reports have already been described in the scientific literature involving free-range or domestic birds. In addition to sharing virulence factors with other enteropathogens, such as *Escherichia coli*, *Hafnia alvei* has been reported to cause serious infections in laying hens [64, 65]. Miniero Davies et al. [66] described an outbreak of mortality associated with *E. tarda* affecting fish, domestic ducks and a wild heron that shared a lake located on a farm in the state of São Paulo, Brazil. Davies et al. [67] described *Klebsiella pneumoniae* expressing virulence and antibiotic resistance genes in psittacine and passerine birds from illegal trade. *Cronobacter sakazakii* has been reported in broilers with clinical signs causing high mortality and decreased egg production [68]. *Proteus* sp. are also potentially pathogenic for birds as the cause of foot injuries and involvement of the respiratory system causing air sacculitis and caseous pneumonia in cases of immunosuppression [69]. Bacteria from the *Arizona* group have often been isolated from feces of adult chickens and turkeys but have also been reported to occur in wild birds, such as the Canadian crane [70, 71]. However, more serious occurrences have been reported in industrial birds, such as mortality in turkeys, as well as clinical signs of Salmonellosis and omphalitis in broiler chickens [72, 1].

It is always more expected to detect cases of antimicrobial resistance in birds raised in captivity than those that live in the wild. In addition to the possibility of inappropriate use of antibiotics, this may occur when birds have greater contact with other animals that possess and disseminate resistant strains [73, 74]. However, our research showed cases of free-living bird strains with relevant antimicrobial resistance rates, mainly involving ampicillin, nalidixic acid and amoxicillin associated with clavulanic acid. Some studies involving free-living birds has also reported resistance to these three antibiotics in isolates of enterobacteria with varying rates. Carreira [75] researched samples of cloacal swabs from free-living birds captured in the Metropolitan region of Fortaleza, Brazil and observed that the acquired resistance rates of amoxicillin associated with clavulanic acid, as well as nalidixic acid, were lower than the results found in this study. Tsubokura et al. [76] analyzed *Escherichia coli* isolates from the feces of several migratory bird species collected in the coastal region of Japan found that less than 10% of the samples were resistant to ampicillin. These same researchers used the feces of 54-day-old Hyline chicks and found the resistance to ampicillin to be approximately 39.0%. These variations can often be attributed to the conditions found in different habitats [77], as demonstrated by several studies that measure resistance levels in isolates from birds under different conditions or captured in different environments [78, 24, 79, 80, 81].

The rate of resistance to meropenem detected in free-living birds in this research should also be highlighted (14.5%). Several studies involving wild birds, free-living or not, as well as domestic birds, present lower rates of resistance to this drug or none at all [82, 83, 30, 84, 85]. However, a more relevant point is the fact that this drug is a high cost carbapenemic with restricted use to hospitals in Brazil. In addition, it is a last resort for the treatment of infections and is widely prescribed to human patients with septic conditions in intensive care of severe infections by Gram-negative hospital pathogens, including Enterobacteriaceae [86, 87, 88]. Although the recommendations for the use of this drug are restricted, the reservoirs of these organisms are increasing, not only in hospitals, but also in the community and the environment. An important new source of resistance development of such organisms is observed in livestock, companion animals and wildlife [89].

Concerning the total number of isolated enterobacterial strains (23.6%) and specifically *Escherichia coli* (15.4%), worrisome rates of multidrug resistance were observed, considering that these are Gram-negative bacteria from free-living animals. Other studies have also demonstrated the occurrence of multidrug resistance of bacteria isolated from cloacal swabs in free-living birds [90, 85]. However, it is not so simple to obtain a proper comparison of data from other studies, since there are few published articles specifically involving free-living birds and isolates of enterobacteria in general from cloacal swabs. One of these studies involves Gray-breasted Parakeets (*Pyrrhura griseipectus*), whose total bacterial isolates presented a lower multidrug resistance rate (11.1%) [30]. Concerning *Escherichia coli*, it is possible to observe that the results in relation to multidrug resistance are the most varied. However, it is possible to find similar rates [15], percentages lower than 5.4% or higher than 23.1% [26, 84].

Densely populated urban areas are historically seen as hotspots for antibiotic resistant bacteria [91, 92] but microorganisms with these characteristics associated with humans have been described in non-clinical environments, such as in remote areas of the planet, far from direct anthropogenic pressure, apparently free from exposure to antibiotics, as in regions of the Amazon, Bolivia and Antarctica. It is suspected that this resistance may have been caused by the existence in these regions of military bases, domestic animals, water, fishing boats, scientific expeditions and/or on-board tourism [93, 94, 95, 96, 97]. Although it is important to emphasize that the cause of antibiotic resistance may not always be related to environmental pressures caused by man, as is the case of those that are naturally induced by microorganisms that produce natural antibiotics [98].

The considerable resistance rates detected in isolates from birds captured in Mulungu, more particularly those tested with ampicillin, amoxicilin+clavulunate, meropenem and nalidixic acid, may indicate that some contact with anthropogenic residues has occurred. Thus, we can consider that wild birds included in our study may be working as indicators of environmental contamination. In this context, we found that free-living birds can be considered victims of the environment in which they live, acquiring multidrug resistant bacteria. At some point, this condition can harm the conservation of species, or they may act as reservoirs of resistant bacteria [97, 99, 100, 101]. Thus, the emergence and evolution of antibiotic resistance among pathogenic bacteria represent a serious public health issue on a global scale [102, 103].

## Conclusion

This study revealed that the investigated wild free-living birds harbor a diverse cloacal microbiota concerning the Enterobaceriaceae family. The phenotypic analysis of the isolates revealed the occurrence of bacterial resistance to several of the antimicrobials tested. Among these, the resistance rates to ampicillin and nalidixic acid can be considered high, since these isolates originated from free-living animals, which naturally suffer a lower selective pressure by antibiotics than domestic ones. The percentage of resistance found to meropenem (14.5%) was also higher than normally expected, since it is a drug with restricted use in hospitals. A relevant multidrug resistance rate was also detected in this study (23.5%), and this shows that birds associated with local extinction risk, such as Ceara Gnateater (*Conopophaga cearae*), are also being affected.

Although this research did not investigate the direct or indirect relationship of wild birds in the Region of Mulungu-CE with sources of contamination, such as sewage water, dumps, crops, soil and domestic or wild animals, it is possible to assume that they could have some contact with contaminating agents, which explains the multidrug resistance rates detected in the cloacal microbiota isolates. Furthermore, birds that have been infected by these microorganisms may also be carrying resistant bacteria to other wild birds or to domestic animals.

## Author Contributions

**Conceptualization:** Antonio Jackson Forte Beleza, Régis Siqueira de Castro Teixeira

**Data curation:** Antonio Jackson Forte Beleza

**Formal analysis:** William Cardoso Maciel, Arianne Silva Carreira

**Funding acquisition:** Antonio Jackson Forte Beleza, Adson Ribeiro Marques, Carlos Henrique Guedes Nogueira

**Investigation:** Régis Siqueira de Castro Teixeira, Antonio Jackson Forte Beleza, Adson Ribeiro Marques, Leandro Rodrigues Ribeiro, Neilton Monteiro Pascoal Filho

**Methodology:** Régis Siqueira de Castro Teixeira, Antonio Jackson Forte Beleza, Bruno Pessoa Lima

**Project administration:** Antonio Jackson Forte Beleza

**Resources:** Antonio Jackson Forte Beleza

**Supervision:** William Cardoso Maciel

**Validation:** William Cardoso Maciel, Isaac Neto Goes da Silva

**Writing ± review & editing:** Ruben Horn Vasconcelos, Régis Siqueira de Castro Teixeira, Antonio Jackson Forte Beleza.

## Acknowledgments

The authors would like to thank the Coordination for the Improvement of Higher Education Personnel (CAPES) for the financial support. We are also grateful for the fundamental logistical support given by Mr. Saulo Rocha from Fazenda Haras Claro, who provided accommodation for the entire team of researchers responsible for field activities.

